# Requirement of ClpX for CtsR dissociation from its operator elements upon heat stress in *Bacillus subtilis*

**DOI:** 10.1101/2025.09.04.674265

**Authors:** Marco Harms, Chelsea Kaden, Larissa M Busch, Vishnu M Dhople, Ulf Gerth, Manuela Gesell Salazar, Stephan Michalik, Zhanetta Zhatarova, Uwe Völker, Alexander Reder

**Affiliations:** University Medicine Greifswald, Center for Functional Genomics of Microbes, Interfaculty Institute for Genetics and Functional Genomics, Greifswald, Germany; University of Greifswald, Center for Functional Genomics of Microbes, Institute of Microbiology, Greifswald, Germany

**Keywords:** *Bacillus subtilis*, class III heat-shock regulon, ClpX, CtsR, ClpE

## Abstract

A sudden increase in temperature triggers *Bacillus subtilis* to activate expression of stress-specific heat shock proteins of the CtsR (**c**lass **t**hree **s**tress gene **r**epressor) regulon to withstand the adverse conditions. Key members of this regulon, such as ATPases, proteolytic subunits and their adaptors, which can assemble to the functional Clp protease system, perform crucial roles in maintaining cellular proteostasis, while their transcription is repressed by CtsR during vegetative growth. Upon heat shock, a conformational change in a thermosensing glycine-rich loop causes CtsR to detach from its DNA operators, enabling the transcriptional activation of the regulon. Novel data from a *clpX*-deficient strain demonstrated that in addition, the presence of the ATPase ClpX is essential for the CtsR dissociation from its DNA binding site. To further elucidate this role of ClpX, we constructed a conditional *clpX* strain, in which *clpX* induction is decoupled from its native transcriptional control. This conditional expression system mimicked a *clpX-*deficient phenotype under non-inducing conditions and restored the wild-type phenotype upon induction. Our results indicate that the full induction of the CtsR regulon, particularly *clpE*, requires both heat and the presence of ClpX, thereby extending the current model for the transcriptional activation of genes repressed by CtsR.

## 1 Introduction

Heat poses a significant threat to the cellular integrity of the Gram-positive bacterium *Bacillus subtilis*, especially in its natural soil habitat, where temperature fluctuations are frequent. Elevated temperatures can disrupt cellular processes by protein misfolding and aggregation, which can severely affect survival. To counteract these adverse effects, *B. subtilis* has evolved an intricate regulatory heat shock stimulon that orchestrates the adaptive heat response (reviewed in (1)). Key regulators of this system include the alternative sigma factor SigB (2) and repressors such as HrcA (3,4) and CtsR (5,6) as well as the response regulator CssR (7,8).

The degradation of misfolded or damaged proteins is crucial for protein homeostasis especially under heat stress conditions. This process is mediated by the gene products of the CtsR regulon, namely key members of the AAA+ superfamily (<u>A</u>TPases <u>a</u>ssociated with various cellular <u>a</u>ctivities). Chaperones and proteases within this family play pivotal roles in maintaining protein quality, while their function relies on ATP hydrolysis (9–11). These bacterial AAA+ complexes typically comprise an unfoldase and a protease subunit, forming hexameric and heptameric rings respectively, that assemble into barrel-like protease complexes (12). The Clp (<u>c</u>aseinolytic <u>p</u>roteases) proteins, integral members of the AAA+ family, function either as chaperones assisting protein refolding (13–15) or in interaction with the proteolytic subunit as protease systems, mediating degradation (16,17). In *B. subtilis*, the ATP-dependent Clp protease proteolytic subunit (ClpP) (18) associates with one of the ATPase subunits ClpC, ClpX or ClpE (19). The specific ATPase subunit dictates substrate specificity, determining which proteins are targeted for degradation (20,21).

During vegetative growth, the dimeric helix-turn-helix (HTH) CtsR repressor protein prevents the transcription of its regulon members by binding to a highly conserved heptanucleotide direct repeat [A/GGTCAAA] within the −35 and −10 core promoter regions (5,22,23). Cooperative binding of CtsR inhibits transcriptional initiation by the RNA polymerase (23). The CtsR regulated genes include the *clpC* operon (*ctsR*-*mcsA*-*mcsB*-*clpC*), as well as the *clpP* and *clpE* gene. Each of these elements is controlled by multiple promoters: the *clpC* operon and *clpP* are preceded by one σ^B^-type and σ^A^-type promoter, while the *clpE* gene is driven by two σ^A^-type promoters. Notably, the number of CtsR binding sites varies between the regulated genes, indicating differences in repression tightness by CtsR (see figure 1) (5,22,24–26). At 37°C, the *clpC operon* and *clpP* genes are expressed only at low basal levels due to CtsR repression and the *clpE* gene even remains completely repressed by five CtsR binding sites. CtsR activity is modulated in a feedback-loop by the gene products of its regulon. McsA, encoded by the second gene of the *clpC operon*, promotes CtsR binding to the operator and repression of target genes by supporting CtsR dimerization (27). Moreover, ClpC is thought to enhance CtsR activity, putatively by assisting CtsR folding or dimerization (5,23). In contrast, McsB is kept inactive by interaction with ClpC (28).

**Figure 1:**
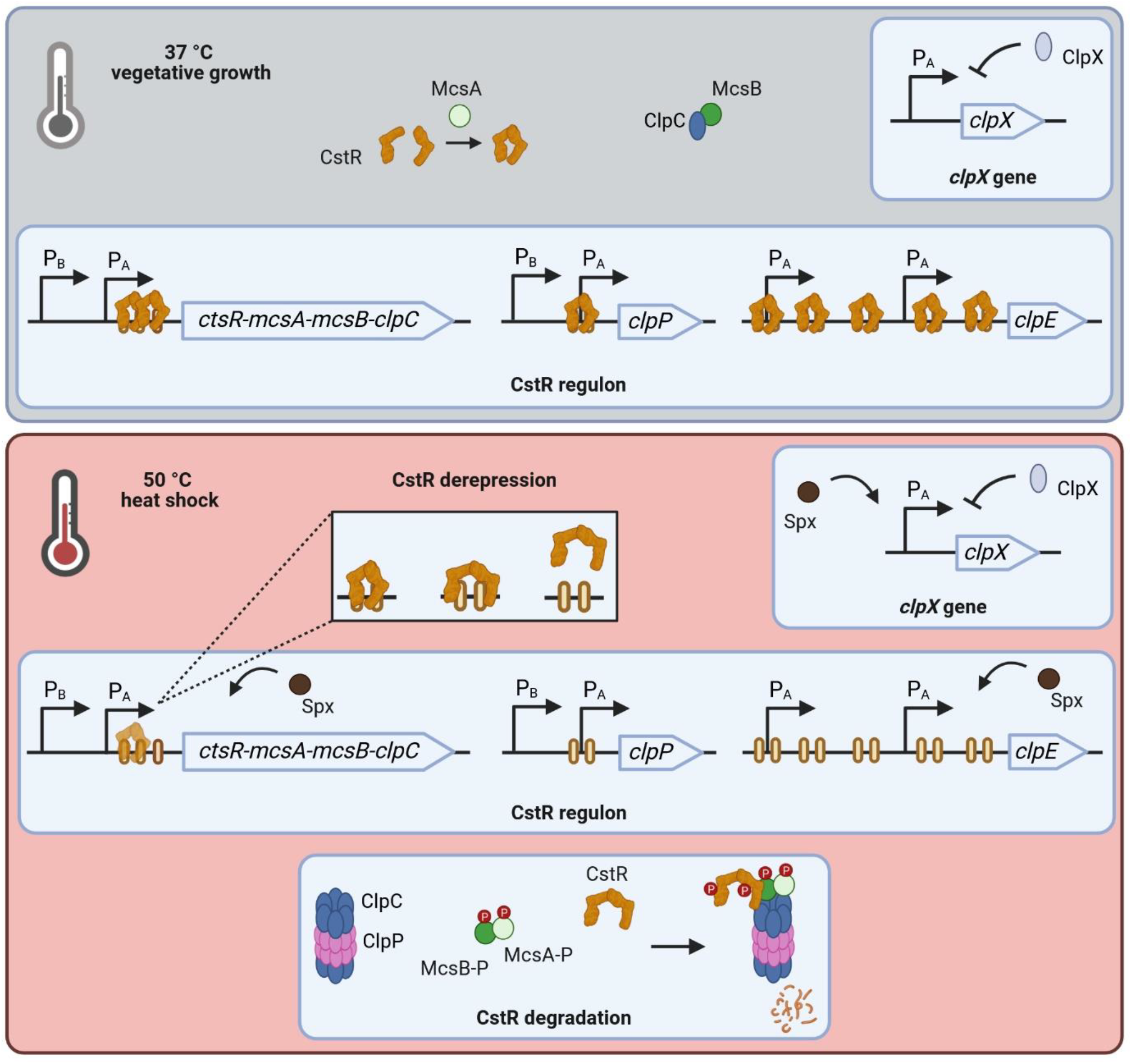
Regulation of *clp* gene expression in B*acillus subtilis*. During vegetative growth CtsR represses transcription of its regulon. McsA putatively assists CtsR dimerization while activity of McsB is abolished by ClpC binding. Transcription of *clpX* as major ATPase is negatively autoregulated. These growth conditions result in expression of low basal level of *clpC* operon members and *clpP*, as well as *clpX*. In contrast, *clpE* is tightly repressed by CtsR. Upon heat stress CtsR changes conformation, which leads to its dissociation from DNA binding sites. Assembly of ClpCP releases McsB. McsA activates McsB leading to phosphorylation of both proteins and subsequent phosphorylation of free, partly unfolded CtsR. The phosphorylated CtsR-McsB-McsA complex is targeted for degradation by ClpCP. As consequence, transcription of now derepressed regulon members is initiated, causing a strong a transient expression. This induction is further accelerated by action of the activator Spx. (Created in BioRender. Völker, U. (2025) https://BioRender.com/h9h78fa)

Sudden temperature upshifts lead to transcriptional activation of the CtsR regulon. In particular, loss of CtsR DNA-binding affinity is triggered by a heat-dependent conformational change of CtsR within the glycine-rich loop adjacent to the helix-turn-helix (HTH) domain (23,29). Subsequent, degradation of unbound CtsR is regulated by a sophisticated two-step mechanism. Formation of the ClpCP protease complex releases the arginine kinase McsB, which is kept inactive by interaction with ClpC at non-heat-stress temperatures (28,30–32). Heat-activated McsA stimulates the protein kinase activity of McsB which results in the phosphorylation of the McsA-McsB complex (28). This complex phosphorylates primarily free CtsR, which is due to the conformational change partly unfolded, on conserved arginine residues (33,34). Phosphorylation labels CtsR for ClpCP-dependent degradation, and simultaneously, phosphorylation in general promotes assembly of the ClpCP complex (27,34,35). Additional degradation of CtsR by ClpEP is described (35). Induction of CtsR regulated genes is transient, leading to elevated transcript and protein level with *clpE* being the most affected due to the number of CtsR binding sites (19,22). Further positive regulation is achieved by the action of Spx on *clpC* operon and the *clpE* gene (36).

Of note, the regulation of the CtsR-independent monocistronic *clpX* gene differs from the other *clp* genes. Like the CtsR-regulated genes, *clpX* is preceded by a σ^A^-type promoter (19,37). However, *clpX* transcription is negatively autoregulated and can be activated by the transcriptional activator Spx under stress conditions (19,36). ClpX functions as the major ATPase at 37°C and is therefore responsible for degradation of CtsR *via* the ClpXP proteolytic complex (23). Although transcription of *clpX* is upregulated upon heat stress, protein levels remain largely unchanged (19,37).

In this study, we report that both the presence of ClpX and heat are prerequisites for the CtsR dissociation of its binding site, thus achieving full induction of the CtsR regulon. Given the experimental data of the newly generated in-frame *clpX* deletional and conditional *clpX* strain, we hypothesize that the chaperone function of ClpX supports the conformational change required for CtsR to detach from its heptanucleotide operator during heat stress.

## 2 Material and Methods

### 2.1 Generation of mutant strains

Genetic modifications were incorporated into the chromosome of *B. subtilis* BSB1 (38). To generate the mutant strains (BZZ01-BZZ05, see table 1), a modified PCR protocol (39) was utilized. This method resulted in generation of linear DNA fragments including the resistance cassette, replacing the target gene and homologous up- and downstream fragments of the desired mutation site. Obtained PCR products were purified and fused according to a procedure described in Dittmar *et al*. (40). Used primers are detailed in supplementary table S1: Primer pairs containing either “up” or “do” are considered primer pairs for the flanking homologous fragments, primer pairs tagged with “Res” were used to amplify the resistance cassette. For the transformation of *B. subtilis* BSB1, modified competence media (MC) according to Smith *et al*. was used (41). 20 ml MC medium was inoculated to OD500 of 0.05 with an exponentially growing overnight LB culture and cultivated at 37 °C, 220 rpm (Innova 4230 Refrigerated Incubator Shaker, New Brunswick). When the MC culture reached an OD500 of 2.0, 2 ml were transferred to another cultivation tube containing 500 µg of respective PCR fragment and further incubated at 90 rpm. After one hour, 1 ml LB medium was added and cultivation continued for 3 hours. The respective mutants were selected on LB agar plates containing either 1 µg/ml erythromycin, 5 µg/ml kanamycin, 5 µg/ml phleomycin, 5µg/ml chloramphenicol or 200 µg/ml spectinomycin. Chromosomal DNA was isolated and mutant clones were verified by Sanger sequencing. The strains BZZ01, BZZ03, BZZ04 and BZZ05 were designed as in-frame deletions, in which the resistance cassette replaces the target at its original chromosomal locus, while in case of strain BZZ02, the phleomycin cassette was integrated with an artificial SigA promoter as well as Shine-Dalgarno sequence. For the conditional *clpX* mutant (BCSH01), the native SigA promoter of the *clpX* gene was replaced to an artificial SigA promoter with two tetracycline operators. Furthermore, *tetR* gene was cloned upstream of the chloramphenicol resistance cassette. Expression of the repressor TetR results in binding to its operators upstream of *clpX* and consequent repression of transcriptional initiation under vegetative growth conditions. Addition of anhydrotetracycline, which sequesters TetR by binding, allowed *clpX* transcription and therefore selective and independent control of the ClpX protein level, regardless of its natural regulation. Detailed information on the construction can be found in figure 4A. BCSH04 (conditional *clpX ΔctsR*) was generated by amplifying a linear DNA fragment of chromosomal DNA of BCSH01 with the primers 1_cond_ClpX_up_for and 6_cond_ClpX_do_rev followed by transformation in the *B. subtilis* strain BZZ05.

**Table 1.** *Bacillus subtilis* strains used in the experiments

| Strain | Relevant features | Reference | Comment |
| --- | --- | --- | --- |
| *B. subtilis* BSB1 | *trp*<sup>+</sup> | (38) |  |
| BZZ01 | *trp*<sup>+</sup>, *clpC*::*km*<sup>R</sup> | This study | Primers in table 2 |
| BZZ02 | *trp*<sup>+</sup>, *clpE*::*phleo*<sup>R</sup> | This study | Primers in table 2 |
| BZZ03 | *trp*<sup>+</sup>, *clpP*::*spec*<sup>R</sup> | This study | Primers in table 2 |
| BZZ04 | *trp*<sup>+</sup>, *clpX*::*erm*<sup>R</sup> | This study | Primers in table 2 |
| BZZ05 | *trp*<sup>+</sup>, *ctsR*::*km*<sup>R</sup> | This study | Primers in table 2 |
| BCSH01 | *trp*<sup>+</sup>, *tig-tetR-cat*<sup>R</sup>-*P*<sub>TRE</sub>-*clpX* | This study | Primers in table 2 |
| BCSH04 | *trp*<sup>+</sup>, *ctsR*::*km*<sup>R</sup>, *trp*<sup>+</sup>, *tig-tetR-cat*<sup>R</sup>-*P*<sub>TRE</sub>-*clpX* | This study | Primers in table 2 |

### 2.2 Cultivation for stress experiments and cell sampling

Cultivations were conducted in at least three biological replicates, in a darkened air incubator at 220 rpm and 37°C (Innova 4230 Refrigerated Incubator Shaker, New Brunswick). A serial dilution was generated by adding 500 µl of the glycerol stock in 5 ml of Lysogeny Broth (LB) medium in disposable reagent tubes (13 ml) and mixed thoroughly. Next, 700 µl was transferred into the second disposable reagent tube, and the process was repeated up to the 12^th^ dilution and incubated for a maximum of 9 h. On the following day, precultures were inoculated with exponentially growing overnight cultures (OD540nm = 0.6 - 1.0) in 30 ml of pre-warmed LB medium to a starting OD540nm of 0.05 and incubated further. As soon as the preculture reached exponential growth, the appropriate volume was transferred to a pre-warmed reaction tube (15 ml), centrifuged at 2400 xg at room temperature (RT) for 1 min. The pellet was resuspended in 100 ml Spizizen’s minimal medium (42) with 0.1 % yeast extract to a starting OD500nm of 0.05 to inoculate the main culture. In the case for the respective condition of BCSH01, 30 min prior to reaching the OD500nm of 0.4 in the main culture, 20 ng/ml anhydrotetracycline (aTc) was added to induce ClpX protein concentrations comparable to those of the wild-type. Once the cultures reached an optical density of OD500 nm = 0.4, 16 OD units were harvested, with 8 OD allocated for RNA and the other 8 OD for protein preparation. Harvest of a defined number of OD units enables sampling of similar cell, RNA and protein content for further processing. Samples were immediately cooled down with liquid nitrogen, centrifuged at 10.000 xg and 4 °C for 3 min. Supernatant was discarded and cell pellets were stored at −70°C. Main culture flasks were immediately transferred into a 50 °C water bath (OLS Aqua Pro, GRANT) and incubated at 150 rpm linear shaking. After 10 min and 30 min, additional samples were collected as described above.

### 2.3 RNA Isolation

RNA-Isolation from the harvested cell pellets was achieved according to Harms *et al*. (43) In brief, cells were mechanically disrupted using the Dismembrator (Retsch) bead mill and RNA was subsequently extracted using the acid phenol method described by Majumdar *et al*. (44).

### 2.4 Quantitative Near-Infrared (NIR) Northern blot analysis

NIR-Northern blots were performed according to the protocol supplied by ProTec Diagnostics GmbH. Transcript sizes were determined using the RNA TRUE Ladder (ProTec Diagnostics). To assess RNA quality and validate equal loading, the blots were stained with methylene blue. A *clpE* RNA probe was biotin-labeled by *in vitro* transcription using the MEGAscript^TM^ T7 Transcription Kit (Invitrogen) and a gene-specific PCR product with a T7 promoter (for primer see supplementary table S1). For fluorescence detection of the Northern blot signal the Odyssey® CLx imaging system (LICORbio Biosciences) was used. For better comparison, ratios of detected signals were calculated by setting the wildtype signal after heat stress (t10) to 100% as reference induction.

### 2.5 Quantitative Near-Infrared (NIR) Western blot analysis

By using an SDS-PAGE (Mini-Protean), 4 µg of the protein lysate were loaded into each lane and separated. Transfer of the separated proteins to a polyvinylidene difluoride (PVDF) membrane was achieved by the protocol of Gerth *et al*. (19). Detection of NIR Western blots was performed following the protocol of LICORbio Biosciences (45), using the Odyssey® CLx imaging system (LICORbio Biosciences). A polyclonal antibody against ClpX (dilution 1:5.000) as described in (19) was used.

### 2.6 Proteome analysis

Protein lysates were generated as described previously (46). Briefly, cell pellets were disrupted using the Dismembrator bead mill (Retsch) followed by nuclease treatment, ultra-sonication and centrifugation. Protein lysates were precipitated according to the protocol described by Nickerson *et al*. (47) and resolubilized in fresh 20 mM HEPES buffer (pH 8.0) with 1% SDS. Precipitation was performed for buffer exchange, since components of the cultivation media would interfere with the protein concentration determination using the Micro BCA™ Protein Assay Kit (Thermo Fisher Scientific) according to Reder *et al*.(48).

For the global label free proteome analyses, the samples of the in-frame mutants were prepared using the SP3 purification and digestion protocol as previously described (49) and the samples of the conditional mutant approach were prepared using the automated SP3 purification and digestion as established in (48) using a modified OT-2 liquid handling robot (Opentrons).

Data independent acquisition (DIA) mass spectrometry (MS) measurements were conducted separately for the in-frame mutant data and for the conditional mutant data.

For the in-frame mutant data, LC-MS/MS analyses were performed using an Orbitrap Exploris™ 480 mass spectrometer in combination with an UltiMate™ 3000 RSLC nano system (both Thermo Fisher Scientific). Detailed specifications of the LC-MS/MS parameters can be found in the supplementary table S2. For the conditional mutant data, measurement was carried out with a Bruker TIMS TOF HT (Bruker Daltonics GmbH) coupled with Evosep One liquid chromatography system using Evosep Pure tips (Evosep Biosystems). For specifications see supplementary table S3.

Data analysis of the DIA experiments were conducted using Spectronaut version 16 (Biognosys AG) and local normalization for the in-frame mutant data. For the conditional mutant data set, data analysis was performed using Spectronaut version 20 and median-median normalized.

Subsequently, the obtained Spectronaut-data was processed using the SpectroPipeR pipeline (50) for quality assessment and MaxLFQ protein level estimation (51). Outlier samples were removed manually from the data sets. Proteins were considered identified if detected with at least two peptides.

For visualization purposes, protein level estimations of mass spectrometry measurements were normalized to the mean of heat stressed wild-type t10 sample protein level estimation. Plots were generated in R v2024.12.1+563 using the tidyverse framework v2.0.0 (52).

## 3 Results

### 3.1 Novel *clpX-*deficient strain reveals unexpected expression patterns of the CtsR regulon

To elucidate the ClpX effect on regulation of the CtsR regulon, the *clpE* gene was selected as the representative marker due to its exclusive regulation by a SigA-type promoter (22), which avoids potential pleiotropic effects associated with SigB. We generated an in-frame *clpX*-deficient strain, which was exposed to heat stress stimulating conditions that activate the CtsR regulon. Using the *clpE* gene as the readout system, we conducted transcriptomic and proteomic profiling with this strain. In agreement with previous studies, no *clpE* transcript was detected either in the *B. subtilis* wild-type (19) or in the novel Δ*clpX* mutant under vegetative growth at 37°C due to the repression by CtsR (see figure 2). Heat stress conditions cause a conformational change within the glycin-rich loop in the dimeric CtsR repressor (29). Ultimately, the dimeric CtsR protein dissociates from its DNA binding and the CtsR regulon is activated. The absence of CtsR already resulted in full induction of *clpE* in the Δ*ctsR* mutant under control conditions. After imposition of heat stress, *clpE* transcript levels increased 1.6-fold resulting in even higher levels compared to the wildtype caused by the activation of the transcriptional regulator Spx. Surprisingly, we observed a decrease in *clpE* transcript levels of approximately 8-fold in the *clpX* mutant strain, when exposed to heat. Given that ClpX is known for its role in protein homeostasis rather than mRNA stability (19,53), we hypothesized that ClpX may indirectly modulate the CtsR regulon during heat stress by affecting the stability or dissociation of CtsR from its DNA binding site.

**Figure 2:**
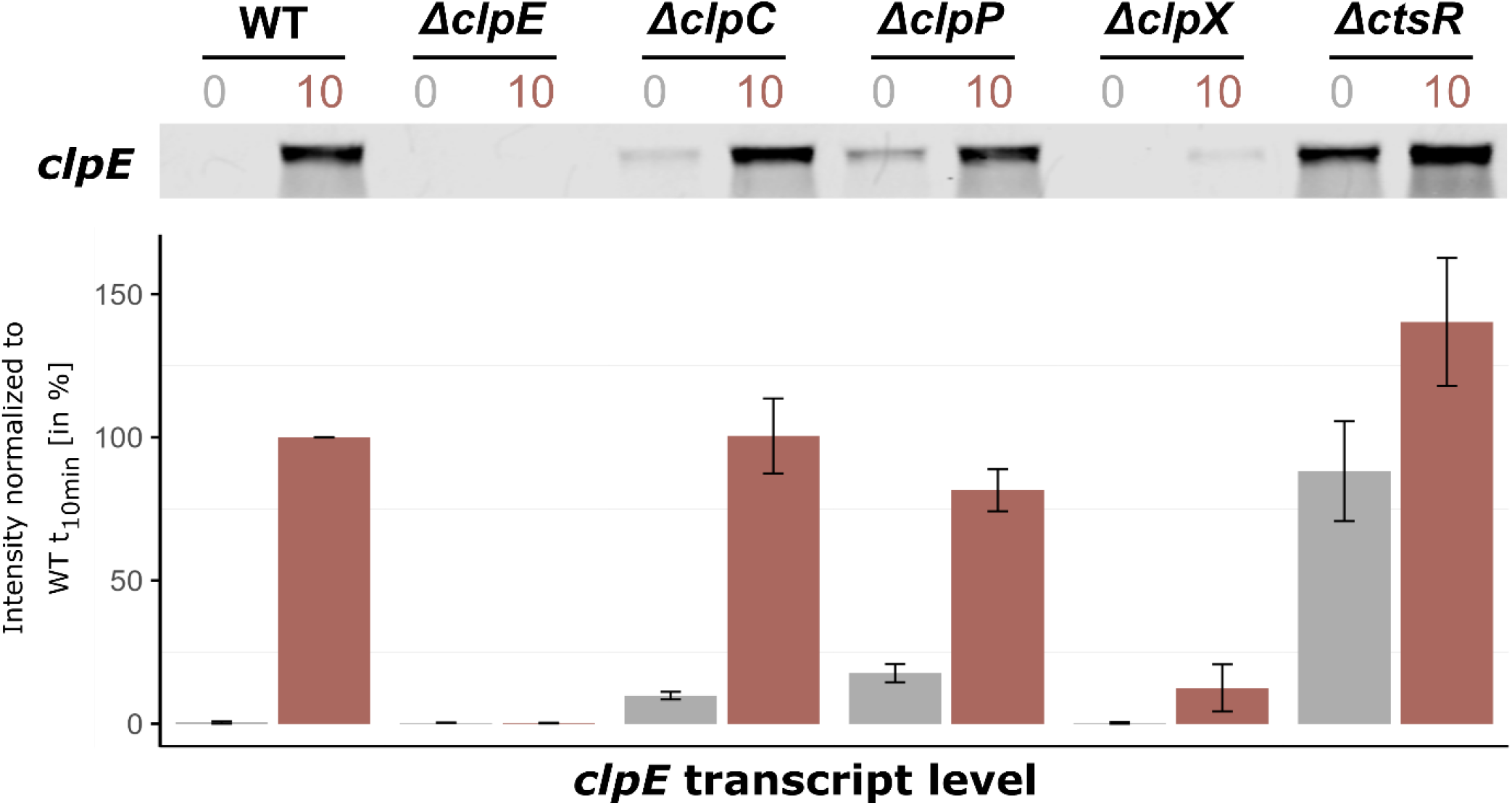
ClpX is involved in derepression of the CtsR regulon under heat stress conditions. Northern blot experiment showing the expression profiles of *clpE*, a gene representative for the CtsR regulon, under control conditions and 10 minutes after heat shock in *B. subtilis* wild-type and its mutants. Cultures were grown in spizizen’s minimal medium with 0.01% yeast extract until the OD_500nm_ of 0.4 (control, t_0_, grey) at 37 °C, then subjected to heat shock in a 50°C water bath (heat stress, t_10_, red). Below, bar plots display the signal quantification of three independent biological replicates, including their standard deviations. The intensities were normalized to the transcript level of the wild-type 10 min after heat shock set to 100%.

### 3.2 Clp protein network dynamics

Having shown that ClpX absence impacts the *clpE* transcript level, we extended our investigation to the *clpC* and *clpP* mutants to clarify whether this observed effect was exclusively mediated by ClpX or whether other Clp proteins were involved. Specifically, we aimed to ascertain if the effects on *clpE* transcript levels were mirrored at protein level and to gain a better understanding of the intricate Clp network in response to heat stress. To achieve this, time-resolved mass spectrometry was performed for a comprehensive proteome monitoring, focusing on the levels of ClpE, ClpC, ClpP, ClpX and CtsR across all previously tested strains (see figure 3). This experimental data revealed that the protein and *clpE* transcript patterns were consistent across all strains, with the exception of the *ctsR* mutant (see figure 2 and 3A). In this case, the lack of CtsR resulted in notably increased levels of ClpC and ClpP (see figure 3B). Elevated concentrations of the ClpCP complex mediated degradation of ClpE, which accounts for the lack of ClpE protein accumulation in the *ctsR* mutant upon heat shock (19).

**Figure 3:**
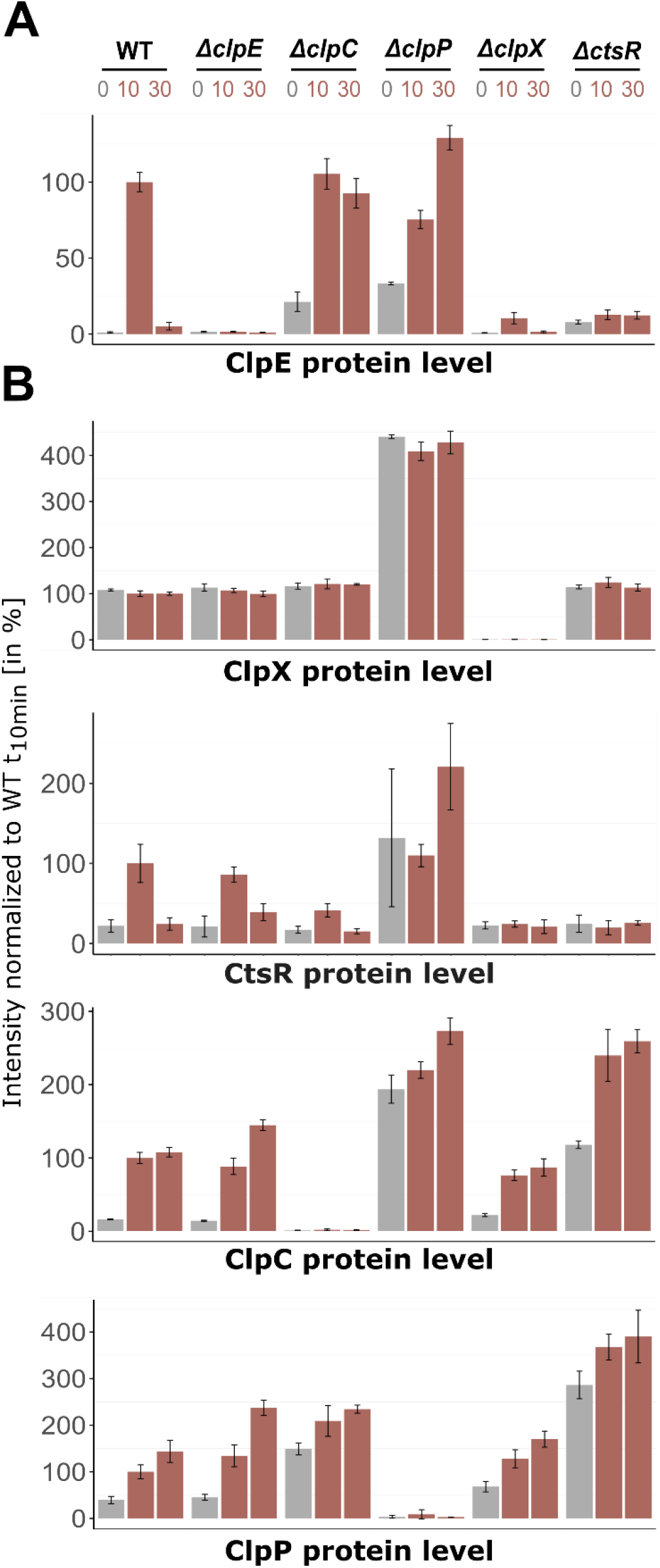
Changes of protein level of the *clpX* gene and CtsR regulon in-frame mutants. Orbitrap Exploris™ 480 mass spectrometer measurements of **(A)** ClpE protein level and **(B)** CtsR regulon members (ClpC, ClpP, CtsR) and ClpX protein level. Relative abundance in percentage normalized to wild-type t_10_ sample are depicted. X-axis titles indicate plotted protein. DIA-MS analysis was performed with four independent biological replicates in different *B. subtilis* strains and their respective standard deviations are displayed. Control conditions (37°C) are highlighted in grey, heat shock conditions (10 and 30 min after 50°C heat shock) in red.

**Figure 4:**
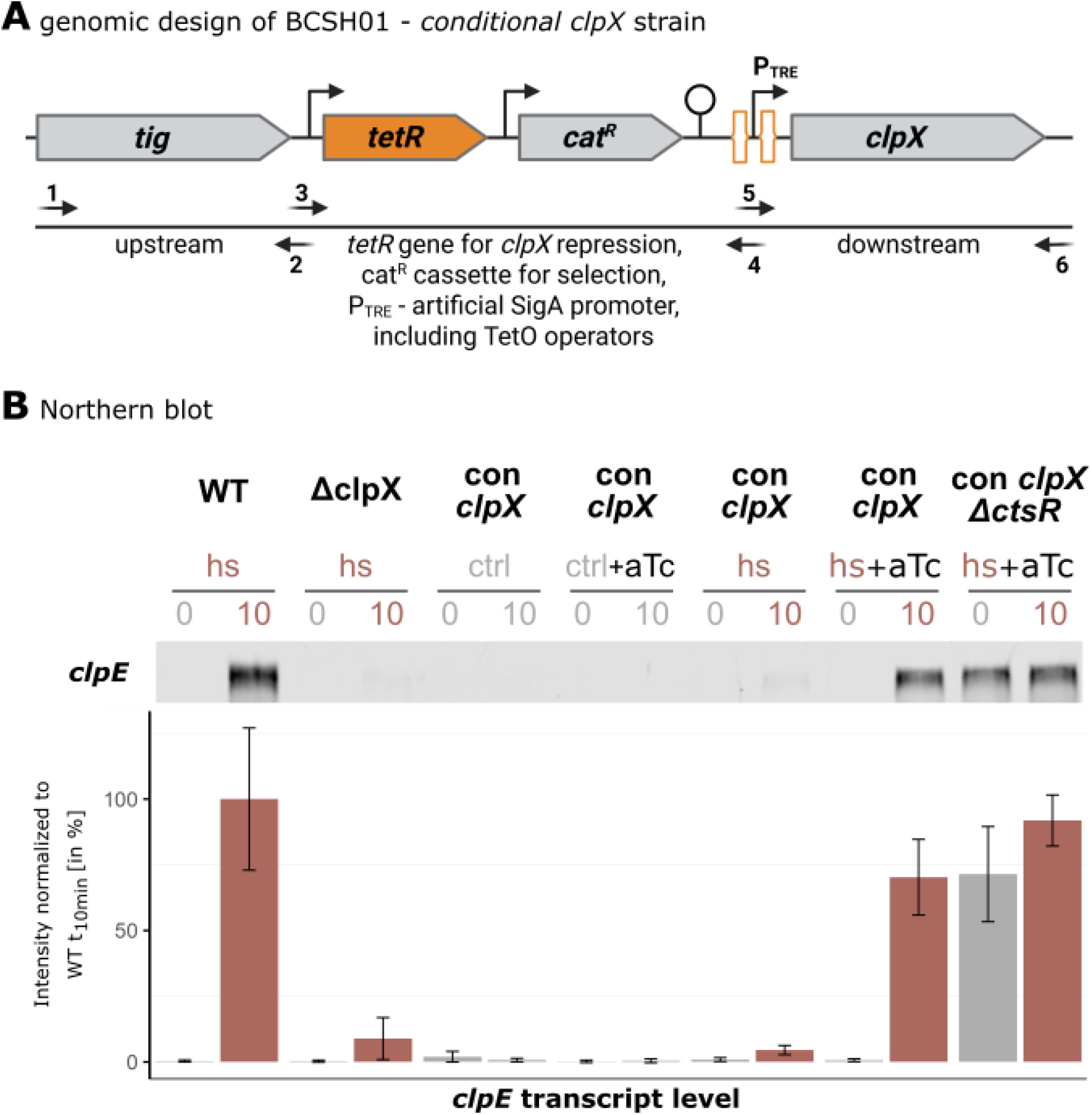
Conditional *clpX* strain construction and Northern blot analysis of *clpE* transcript. **(A)** Detailed genetic construction of the BCSH01 strain. The gene construct containing *tetR, cat*^*R*^ and the artificial P_TRE_ promoter was amplified with the primer pair TetR_PsigA-TRE_for and TetR_PsigA-TRE_rev using the chromosomal DNA of BAR610 as template (2). Primers used to assemble the transformation construct are listed in table 2. TetR repressor binding sites are schematically highlighted in orange. By introducing a strong terminator after the *tetR*-*cat*^*R*^ fusion, transcriptional initiation of *clpX* in the absence of anhydrotetracycline is prevented. (Created in BioRender. Völker, U. (2025) https://BioRender.com/rkxfmk5) **(B)** Representative Northern blot analysis of *clpE* transcript levels in a *B. subtilis* wild-type, its isogenic *clpX* mutant and the conditional *clpX* mutant (BCSH01) under various conditions. Application of heat shock (hs, 50°C in red), vegetative temperatures (ctrl, 37°C in grey) and *clpX* induction 30 min prior to sampling (aTc) is indicated above. Below, bar plots display signal quantification of three independent biological replicates, including their standard deviations normalized to wild-type *clpE* levels 10 min after ongoing heat shock (set as 100%).

When compared to the *B. subtilis* wild-type, we found that the ClpX-deficient strain showed only slightly increased ClpE levels in response to heat stress, which were 9.6-fold lower (see figure 3A). This pattern resembled the data obtained for *clpE* on transcript level. Interestingly, the ClpX effect was also extended to CtsR, as the *clpX*-deficient strain possessed lower and non-inducible CtsR levels in comparison to the wild-type (figure 3B). These findings implied that the presence of ClpX is already essential in the transcriptional derepression of the CtsR regulon. In particular, the appropriate dissociation of CtsR from its DNA-binding motif under heat stress might be compromised in the absence of ClpX.

In the *clpC* mutant, the elevated basal levels of the *clpE* transcript lead to a corresponding increase in ClpE protein levels (see figure 2 and 3A). Derré *et al*. postulated a positive role of ClpC on CtsR activity by assisting correct folding and subsequent repression or protecting it from degradation by ClpXP (23). Wild-type-like CtsR level in the *clpC* mutant (see figure 3B) supported the notion that lack of ClpC may lead to inactive CtsR, resulting in the observed increase in *clpE* transcript levels under non-inducing conditions. Reduced repression by CtsR impacted not only *clpE* but also other regulon members, such as ClpP (see figure 3B). Still, the overall regulation remained distinct from what we have seen in the *clpX*-deficient strain. Similarly, while the *clpP* mutant exhibited accumulation of all proteins of interest in this study due to the loss of the essential proteolytic subunit, the transcriptional and proteomic patterns observed in the absence of ClpX were completely different. Thus, the observed particular effect is confined to the lack of ClpX only, highlighting its unique role in modulating the dissociation of CtsR from its DNA binding sites and thereby regulating the broader CtsR regulon under stress conditions.

### 3.3 ClpX affects derepression of the CtsR regulon

In the next step, we generated a conditional mutant (con *clpX*) strain to precisely control the *clpX* expression levels by addition of anhydrotetracycline (aTc), thereby decoupling *clpX* expression from its natural regulatory stimuli (see figure 4A). This approach allowed us to dissect the direct role of ClpX in real-time, while simultaneously eliminating heat-associated pleiotropic effects. To ensure accurate comparisons, we first titrated the aTc concentration to match the ClpX expression levels observed in *B. subtilis* wild-type (see supplementary figure S1). This allowed examination of the ClpX’s effect under finely tuned conditions to directly correlate changes in the expression of the CtsR regulon.

Integration of the data of *clpE* transcript and protein levels revealed coherent behavior of the investigated strains (see figure 4B and 5A). As expected, in the absence of heat, no induction was observed for the *clpE* gene (see figure 4B) or any corresponding production of ClpE protein (see figure 5A). Without the heat shock, ClpX alone is insufficient to initiate derepression of the *clpE* gene. This finding underscored that heat is a critical factor required to trigger the conformational shift in CtsR and its concurrent dissociation from the DNA binding site (29). Notably, the uninduced conditional *clpX* strain exhibited a transcriptional and proteomic profile under heat stress that was almost identical to that of the *clpX* mutant strain. This similarity confirmed that the aTc-induction system effectively mimicked the *clpX* knockout condition, thus validating the conditional *clpX* strain for further investigation of the ClpX-dependent regulatory mechanism. When ClpX was induced to levels comparable to those of the wild-type and heat stress was applied, the transcriptional and proteomic profiles resembled those of the *B. subtilis* wild-type strain with only 1.2-fold difference (see figure 4B and 5B). When deleting *ctsR* and inducing ClpX production, *clpE* transcription was upregulated even under control (37°C) conditions. However, ClpE protein level displayed similar intensities as in *clpX* deficient strains. This strong discrepancy may be caused by dysregulation and consequently strong upregulation and accumulation of ClpC and ClpP due to the loss of CtsR (see figure 5B). ClpCP has been reported as the proteolytic complex responsible for the majority of protein degradation (24), likely degrading ClpE to low levels.

**Figure 5:**
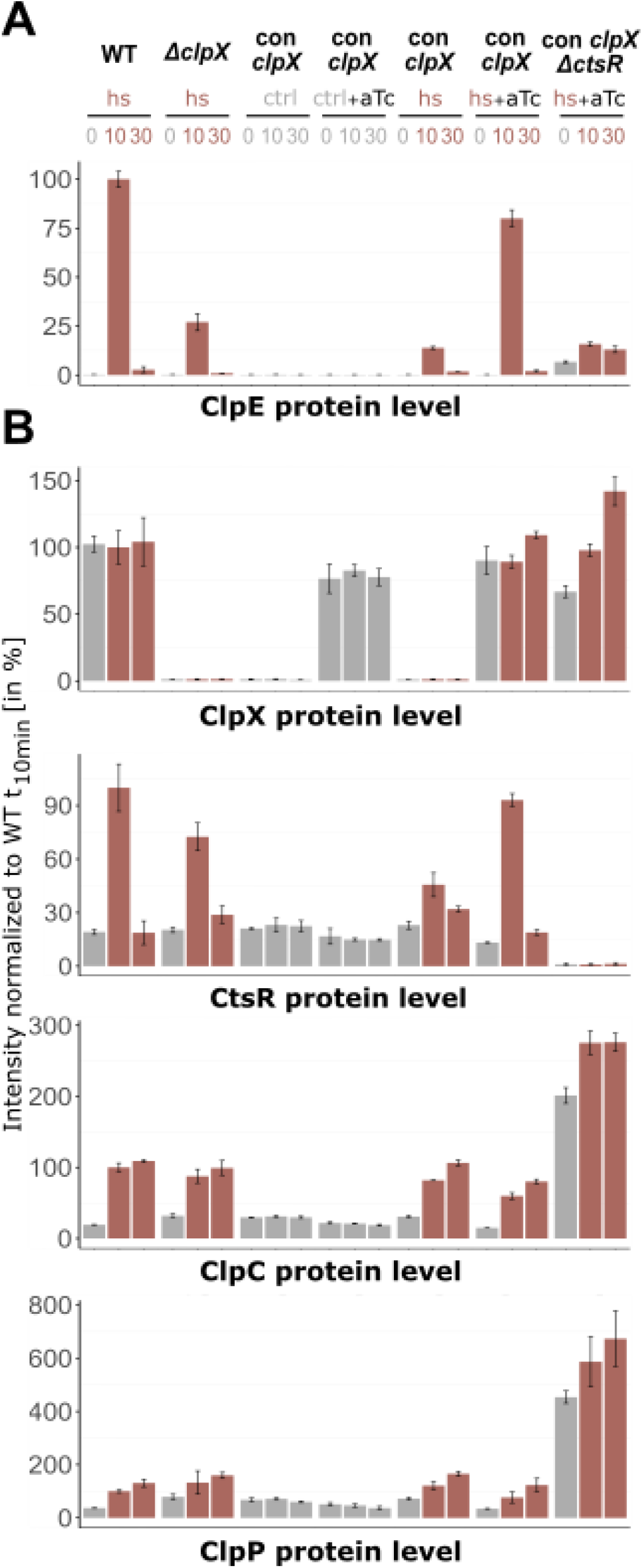
Protein levels in wild-type, *ΔclpX* and conditional *clpX* strains under non-inducing and inducing conditions. Bruker TIMS TOF HT measurement of **(A)** ClpE protein level and **(B)** CtsR regulon members (ClpC, ClpP, CtsR) and ClpX protein level. Relative abundance in percentage normalized to heat stressed wild-type t_10_ sample are depicted. X-axis titles indicate plotted protein. Application of heat shock (hs), vegetative growth (ctrl) and induction (aTc) is indicated above. DIA-MS analysis was performed with three independent biological replicates in different *B. subtilis* strains and their respective standard deviations are displayed. Control conditions (37°C) are highlighted in grey, heat shock conditions (10 and 30 min after 50°C heat shock) in red.

## 4 Discussion

In this study, we elucidated the role of ClpX in the regulation of the CtsR regulon in *B. subtilis* under heat stress conditions. Heat and the presence of ClpX are both prerequisites for derepression of the CtsR regulon, a function of ClpX that extends beyond its established role in protein homeostasis. According to current knowledge, CtsR is regarded as a dimeric DNA-binding protein that binds to a conserved heptanucleotide direct repeat sequence (A/GGTCAAA NAN A/GGTCAAA) overlapping with the transcription initiation site or the −35 and −10 boxes of the CtsR regulon promoter and thereby effectively preventing the transcription of its target genes (5,22). CtsR interacts with the major groove of DNA *via* its β-hairpin structure and with the minor groove *via* its helix-turn-helix (HTH) domain (33). The ability of CtsR to sense temperature changes is attributed to its glycine-rich loop which comprises the residues RGGGGY, located at positions G64-G67 adjacent to the HTH domain (29,33). Our congruent transcript and protein patterns for the *clpE* gene under heat stress conditions demonstrated that ClpX likely performs a crucial chaperone function for the proper dissociation of CtsR.

In contrast to a study conducted by Gerth and coworkers 20 years ago *clpE* mRNA levels in a *clpX* strain indicated very similar expression patterns to a *B. subtilis* wild-type under control (37°C) as well as heat shock (50°C) conditions (19). Resequencing of this *clpX*-deficient strain (BEK90) revealed that this mutant retains part of a N-terminal zinc finger motif, native *clpX* promoter architecture as well as original protein translation sequences (Shine-Dalgarno sequence and ATG start codon) (see supplementary figure S2) (37,54). It has been shown that, in *E. coli*, a zinc finger-like domain of the heat-shock chaperone DnaJ is essential for the recognition and binding to proteins (55). Assuming similar properties for ClpX, it is possible that the remaining translated protein still exhibits residual function. Sequencing of the here constructed in-frame *clpX* mutant strain, as well as congruent data from the uninduced conditional *clpX* strain, verify the complete lack of ClpX.

Our data indicate that upon heat stress conditions, ClpX seems to influence the heat-induced conformational change of CtsR and the induction of the CtsR regulon. The chaperone function of ClpX may be involved in stabilizing an intermediate conformation of CtsR or directly assisting in the refolding of CtsR into a non-DNA-binding-state. Experimental data from Δ*clpC* and Δ*clpP* mutants have shown that this phenomenon is solely attributed to ClpX (see figure 3). Based on these new insights, we propose an adjusted revised model of CtsR regulation that builds on the work of Elsholz and coworkers (see figure 6) (29).

**Figure 6:**
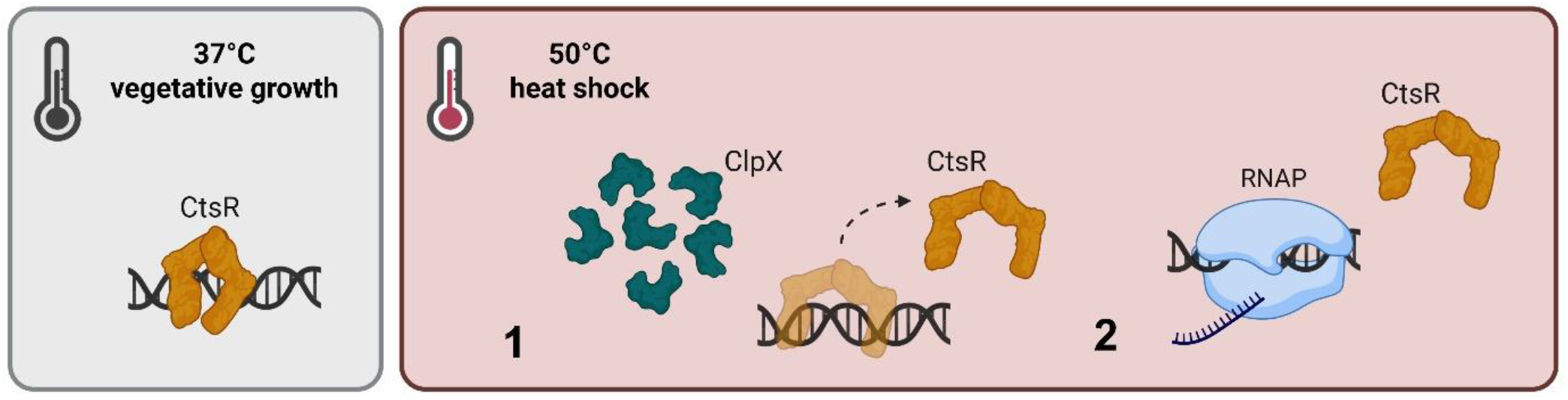
Adjusted model for heat- and ClpX-dependent regulation of the CtsR activity. CtsR operates as a dimeric DNA repressor under control conditions and binds to its conserved heptanucleotide sequence, preventing transcription of the CtsR regulon. (1) A sudden increase in temperature, results in a heat-induced conformational change of CtsR, mediated by the chaperone function of ClpX. Thus, transcriptional initiation of regulated genes is activated (2). (Created in BioRender. Völker, U. (2025) https://BioRender.com/6gz6yh3)

CtsR represses its regulon by binding to its conserved DNA binding motif under vegetative growth conditions. Simultaneously, McsB is kept inactive by the complex formation with ClpC (28). Heat exposure results in a conformational change within the winged HTH-domain of CtsR, a process likely supported by the chaperone function of ClpX. This conformational state of CtsR loses its DNA binding affinity and detaches from its DNA binding site, while the transcription of the CtsR regulon is initiated. Non-functional CtsR is targeted for phosphorylation by the heat-activated McsB-P/McsA complex and thereby tagged for ClpCP-dependent degradation (32).

The current model does not clearly describe how CtsR is reactivated as a DNA-binding repressor under heat stress conditions. According to the model, CtsR reactivation occurs despite ongoing heat stress and the potential heat-induced conformational change of newly synthesized CtsR. We suggest that the availability of ClpX during heat stress plays a pivotal role in the feedback loop of the CtsR regulon. Our data show that ClpX protein levels under heat stress were identical to those observed during vegetative growth. A study by Kirstein *et al*. revealed that Clp-associated proteins primarily localize in clusters at the cell poles or mid-cell region under heat stress to recycle damaged proteins (56). In case, a significant portion of ClpX is sequestered in a complex with ClpP, it may no longer support the conformational shift required for CtsR to effectively dissociate from DNA. Thus, we assume ClpX availability is critical for maintaining the balance of CtsR activity and for regulating CtsR-dependent genes under heat-stress conditions.

## Supporting information

Combined Supplemental Data

## Data availability statement

The mass spectrometry proteomics data for both measurements (in-frame mutants and conditional mutants) as well as the protein databases have been deposited to the ProteomeXchange Consortium *via* the PRIDE partner repository with the data set identifier MSV000099009 (in-frame mutant data, incl. additional data of *mcsB* mutant) and MSV000099010 (conditional mutant data).

Strains generated in this study are available from the corresponding authors upon request.

## Author Contributions

Marco Harms: Conceptualization, Formal Analysis, Investigation, Validation, Visualization, Methodology, Writing – original draft, Writing – review & editing.

Chelsea Kaden: Formal Analysis, Investigation, Methodology, Visualization, Writing – original draft, Writing – review & editing.

Stephan Michalik: Data curation, Formal Analysis, Validation, Software, Writing – review & editing.

Larissa Milena Busch: Data curation, Formal Analysis, Writing – review & editing.

Ulf Gerth: Conceptualization, Supervision, Writing - review & editing.

Manuela Gesell Salazar: Validation, Writing - review & editing.

Vishnu M Dhople: Validation, Writing - review & editing.

Zhanetta Zhatarova: Investigation, Writing - review & editing.

Marc Schaffer: Investigation, Writing - review & editing.

Uwe Völker: Conceptualization, Funding Acquisition, Project Administration, Supervision, Writing - review & editing.

Alexander Reder: Conceptualization, Validation, Supervision, Writing - original draft, Writing - review & editing.

## Acknowledgements

We appreciate the support from ProTec Diagnostics providing the quantitative fluorescence-based near-infrared (NIR) Northern blot protocol. Schematic illustrations were created using BioRender and Inkscape.

## References

1. Schumann W. The Bacillus subtilis heat shock stimulon. Cell Stress Chaperones. 2003;8(3):207.

2. Price CW, Fawcett P, Cérémonie H, Su N, Murphy CK, Youngman P. Genome-wide analysis of the general stress response in Bacillus subtilis. Mol Microbiol. 2001 Aug;41(4):757–74.

3. Schulz A, Schumann W. hrcA, the first gene of the Bacillus subtilis dnaK operon encodes a negative regulator of class I heat shock genes. J Bacteriol. 1996 Feb;178(4):1088–93.

4. Yuan G, Wong SL. Isolation and characterization of Bacillus subtilis groE regulatory mutants: evidence for orf39 in the dnaK operon as a repressor gene in regulating the expression of both groE and dnaK. J Bacteriol. 1995 Nov;177(22):6462–8.

5. Derré I, Rapoport G, Msadek T. CtsR, a novel regulator of stress and heat shock response, controls clp and molecular chaperone gene expression in Gram-positive bacteria. Mol Microbiol. 1999 Jan;31(1):117–31.

6. Krüger E, Hecker M. The First Gene of the Bacillus subtilis clpC Operon, ctsR, Encodes a Negative Regulator of Its Own Operon and Other Class III Heat Shock Genes. J Bacteriol. 1998 Dec 15;180(24):6681–8.

7. Darmon E, Noone D, Masson A, Bron S, Kuipers OP, Devine KM, et al. A Novel Class of Heat and Secretion Stress-Responsive Genes Is Controlled by the Autoregulated CssRS Two-Component System of Bacillus subtilis. J Bacteriol. 2002 Oct 15;184(20):5661–71.

8. Hyyryläinen H, Bolhuis A, Darmon E, Muukkonen L, Koski P, Vitikainen M, et al. A novel two-component regulatory system in Bacillus subtilis for the survival of severe secretion stress. Mol Microbiol. 2001 Sep;41(5):1159–72.

9. Erzberger JP, Berger JM. Evolutionary relationships and structural mechanisms of AAA+ proteins. Annu Rev Biophys Biomol Struct. 2006 Jun 1;35(1):93–114.

10. Gottesman S, Maurizi MR, Wickner S. Regulatory Subunits of Energy-Dependent Proteases. Cell. 1997 Nov;91(4):435–8.

11. Pak M, Hoskins JR, Singh SK, Maurizi MR, Wickner S. Concurrent Chaperone and Protease Activities of ClpAP and the Requirement for the N-terminal ClpA ATP Binding Site for Chaperone Activity. J Biol Chem. 1999 Jul;274(27):19316–22.

12. Martin A, Baker TA, Sauer RT. Rebuilt AAA + motors reveal operating principles for ATP-fuelled machines. Nature. 2005 Oct;437(7062):1115–20.

13. Gottesman S, Wickner S, Maurizi MR. Protein quality control: triage by chaperones and proteases. Genes Dev. 1997 Apr 1;11(7):815–23.

14. Schirmer EC, Glover JR, Singer MA, Lindquist S. HSP100/Clp proteins: a common mechanism explains diverse functions. Trends Biochem Sci. 1996 Aug;21(8):289–96.

15. Wickner S, Maurizi MR, Gottesman S. Posttranslational Quality Control: Folding, Refolding, and Degrading Proteins. Science. 1999 Dec 3;286(5446):1888–93.

16. Hinnerwisch J, Reid BG, Fenton WA, Horwich AL. Roles of the N-domains of the ClpA Unfoldase in Binding Substrate Proteins and in Stable Complex Formation with the ClpP Protease. J Biol Chem. 2005 Dec;280(49):40838–44.

17. Weber-Ban EU, Reid BG, Miranker AD, Horwich AL. Global unfolding of a substrate protein by the Hsp100 chaperone ClpA. Nature. 1999 Sep;401(6748):90–3.

18. Kock H, Gerth U, Hecker M. The ClpP Peptidase Is the Major Determinant of Bulk Protein Turnover in Bacillus subtilis. J Bacteriol. 2004 Sep;186(17):5856–64.

19. Gerth U, Kirstein J, Mostertz J, Waldminghaus T, Miethke M, Kock H, et al. Fine-Tuning in Regulation of Clp Protein Content in Bacillus subtilis. J Bacteriol. 2004 Jan;186(1):179–91.

20. Hoskins JR, Pak M, Maurizi MR, Wickner S. The role of the ClpA chaperone in proteolysis by ClpAP. Proc Natl Acad Sci. 1998 Oct 13;95(21):12135–40.

21. Levchenko I, Smith CK, Walsh NP, Sauer RT, Baker TA. PDZ-like Domains Mediate Binding Specificity in the Clp/Hsp100 Family of Chaperones and Protease Regulatory Subunits. Cell. 1997 Dec;91(7):939–47.

22. Derré I, Rapoport G, Devine K, Rose M, Msadek T. ClpE, a novel type of HSP100 ATPase, is part of the CtsR heat shock regulon of Bacillus subtilis. Mol Microbiol. 1999 May;32(3):581–93.

23. Derré I, Rapoport G, Msadek T. The CtsR regulator of stress response is active as a dimer and specifically degraded in vivo at 37°C. Mol Microbiol. 2000 Oct;38(2):335–47.

24. Krüger E, Witt E, Ohlmeier S, Hanschke R, Hecker M. The Clp Proteases of Bacillus subtilis Are Directly Involved in Degradation of Misfolded Proteins. J Bacteriol. 2000 Jun;182(11):3259–65.

25. Krüger E, Msadek T, Hecker M. Alternate promoters direct stress-induced transcription of the Bacillus subtilis clpC operon. Mol Microbiol. 1996 May;20(4):713–23.

26. Gerth U, Krüger E, Derré I, Msadek T, Hecker M. Stress induction of the Bacillus subtilis clpP gene encoding a homologue of the proteolytic component of the Clp protease and the involvement of ClpP and ClpX in stress tolerance. Mol Microbiol. 1998 May;28(4):787–802.

27. Krüger E, Zühlke D, Witt E, Ludwig H, Hecker M. Clp-mediated proteolysis in Gram-positive bacteria is autoregulated by the stability of a repressor. EMBO J. 2001 Feb 15;20(4):852–63.

28. Kirstein J, Zühlke D, Gerth U, Turgay K, Hecker M. A tyrosine kinase and its activator control the activity of the CtsR heat shock repressor in B. subtilis. EMBO J. 2005 Oct 5;24(19):3435–45.

29. Elsholz AKW, Michalik S, Zühlke D, Hecker M, Gerth U. CtsR, the Gram-positive master regulator of protein quality control, feels the heat. EMBO J. 2010 Nov 3;29(21):3621–9.

30. Kirstein J, Dougan DA, Gerth U, Hecker M, Turgay K. The tyrosine kinase McsB is a regulated adaptor protein for ClpCP. EMBO J. 2007 Apr 18;26(8):2061–70.

31. Lu K, Luo B, Tao X, Luo Y, Ao M, Zheng B, et al. Complex structure and activation mechanism of arginine kinase McsB by McsA. Nat Chem Biol. 2025 Mar;21(3):402–11.

32. Elsholz AKW, Hempel K, Michalik S, Gronau K, Becher D, Hecker M, et al. Activity Control of the ClpC Adaptor McsB in Bacillus subtilis. J Bacteriol. 2011 Aug;193(15):3887–93.

33. Fuhrmann J, Schmidt A, Spiess S, Lehner A, Turgay K, Mechtler K, et al. McsB Is a Protein Arginine Kinase That Phosphorylates and Inhibits the Heat-Shock Regulator CtsR. Science. 2009 Jun 5;324(5932):1323–7.

34. Trentini DB, Suskiewicz MJ, Heuck A, Kurzbauer R, Deszcz L, Mechtler K, et al. Arginine phosphorylation marks proteins for degradation by a Clp protease. Nature. 2016 Nov 3;539(7627):48–53.

35. Miethke M, Hecker M, Gerth U. Involvement of Bacillus subtilis ClpE in CtsR Degradation and Protein Quality Control. J Bacteriol. 2006 Jul;188(13):4610–9.

36. Rochat T, Nicolas P, Delumeau O, Rabatinová A, Korelusová J, Leduc A, et al. Genome-wide identification of genes directly regulated by the pleiotropic transcription factor Spx in Bacillus subtilis. Nucleic Acids Res. 2012 Oct;40(19):9571–83.

37. Gerth U, Wipat A, Harwood CR, Carter N, Emmerson PT, Hecker M. Sequence and transcriptional analysis of clpX, a class-III heat-shock gene of Bacillus subtilis. Gene. 1996 Nov;181(1–2):77–83.

38. Nicolas P, Mäder U, Dervyn E, Rochat T, Leduc A, Pigeonneau N, et al. Condition-Dependent Transcriptome Reveals High-Level Regulatory Architecture in Bacillus subtilis. Science. 2012 Mar 2;335(6072):1103–6.

39. Wach A. PCR-synthesis of marker cassettes with long flanking homology regions for gene disruptions in S. cerevisiae. Yeast. 1996 Mar 15;12(3):259–65.

40. Dittmar D, Reder A, Schlüter R, Riedel K, Hecker M, Gerth U. Complementation studies with human ClpP in Bacillus subtilis. Biochim Biophys Acta BBA - Mol Cell Res. 2020 Sep;1867(9):118744.

41. Smith DR, Kearns DB, Burton BM. ComI inhibits transformation in Bacillus subtilis by selectively killing competent cells. Henkin TM, editor. J Bacteriol. 2024 Jul 25;206(7):e00413–23.

42. Harwood CR, Cutting SM. Molecular biological methods for Bacillus. Chichester New York Brisbane [etc.]: J. Wiley & sons; 1990. (Modern microbiological methods).

43. Harms M, Wolfgramm H, Schedlowski M, Michalik S, Hildebrandt P, Schaffer M, et al. Characterization of the MgsR-dependent promoter structure in Bacillus subtilis —application of a novel pHIS plasmid-based screening system for promoter element analysis. Nucleic Acids Res. 2025 Jul 8;53(13):gkaf636.

44. Majumdar D, Avissar YJ, Wyche JH. Simultaneous and rapid isolation of bacterial and eukaryotic DNA and RNA: a new approach for isolating DNA. BioTechniques. 1991 Jul;11(1):94–101.

45. Near-Infrared Western Blot Detection Protocol [Internet]. [cited 2025 Aug 28]. Available from: https://www.licorbio.com/support/contents/applications/western-blots/fluorescent-western-blot-detection-protocol.html?Highlight=on-cell%20western%20assay.

46. Harms M, Michalik S, Hildebrandt P, Schaffer M, Gesell Salazar M, Gerth U, et al. Activation of the general stress response sigma factor SigB prevents competence development in Bacillus subtilis. Msadek T, editor. mBio. 2024 Dec 11;15(12):e02274–24.

47. Nickerson JL, Doucette AA. Rapid and Quantitative Protein Precipitation for Proteome Analysis by Mass Spectrometry. J Proteome Res. 2020 May 1;19(5):2035–42.

48. Reder A, Hentschker C, Steil L, Gesell Salazar M, Hammer E, Dhople VM, et al. MassSpecPreppy— An end-to-end solution for automated protein concentration determination and flexible sample digestion for proteomics applications. PROTEOMICS. 2024 May;24(9):2300294.

49. Ganske A, Busch LM, Hentschker C, Reder A, Michalik S, Surmann K, et al. Exploring the targetome of IsrR, an iron-regulated sRNA controlling the synthesis of iron-containing proteins in Staphylococcus aureus. Front Microbiol. 2024 Jul 5;15:1439352.

50. Michalik S, Hammer E, Steil L, Salazar MG, Hentschker C, Surmann K, et al. SpectroPipeR—a streamlining post Spectronaut® DIA-MS data analysis R package. Nikolski M, editor. Bioinformatics. 2025 Mar 4;41(3):btaf086.

51. Cox J, Hein MY, Luber CA, Paron I, Nagaraj N, Mann M. Accurate Proteome-wide Label-free Quantification by Delayed Normalization and Maximal Peptide Ratio Extraction, Termed MaxLFQ. Mol Cell Proteomics. 2014 Sep;13(9):2513–26.

52. Wickham H, Averick M, Bryan J, Chang W, McGowan L, François R, et al. Welcome to the Tidyverse. J Open Source Softw. 2019 Nov 21;4(43):1686.

53. Gerth U, Kock H, Kusters I, Michalik S, Switzer RL, Hecker M. Clp-Dependent Proteolysis Down-Regulates Central Metabolic Pathways in Glucose-Starved Bacillus subtilis. J Bacteriol. 2008 Jan;190(1):321–31.

54. Gottesman S, Clark WP, De Crecy-Lagard V, Maurizi MR. ClpX, an alternative subunit for the ATP-dependent Clp protease of Escherichia coli. Sequence and in vivo activities. J Biol Chem. 1993 Oct;268(30):22618–26.

55. Szabo A, Korszun R, Hartl FU, Flanagan J. A zinc finger-like domain of the molecular chaperone DnaJ is involved in binding to denatured protein substrates. EMBO J. 1996 Jan 15;15(2):408–17.

56. Kirstein J, Strahl H, Molière N, Hamoen LW, Turgay K. Localization of general and regulatory proteolysis in Bacillus subtilis cells. Mol Microbiol. 2008 Nov;70(3):682–94.

