## Supplementary material for "Requirement of ClpX for CtsR dissociation from its operator elements upon heat stress in *Bacillus subtilis*": Combined Supplemental Data

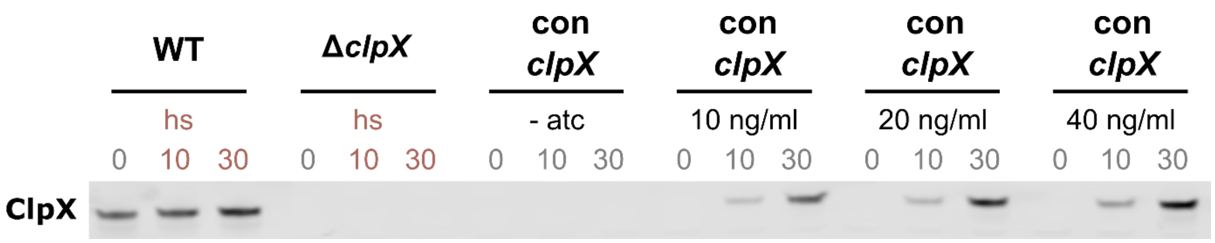


**Supplementary Figure S1: Determination of optimal aTc concentration used for conditional *clpX* induction obtaining wild-type protein levels.**

Western blot analysis ClpX protein levels in a *B. subtilis* wild-type, its isogenic *clpX* mutant and the conditional *clpX* mutant (con *clpX* = BCSH01) under various conditions: 10 min and 30 min after heat shock (50°, highlighted in red) and the addition of 0, 10, 20 or 40 ng/ml anhydrotetracycline 30 min prior to the sampling (atc). Grey indicates growth at 37°C. Based on this experimental result, the addition of 20 ng/ml aTc 30 min prior to the sampling was chosen as the concentration for the induction of the *clpX* gene to ensure ClpX levels comparable to that of the wild-type.

```
                DNA alignment

       -35 Box                       -10 Box
WT     GTTACCGTCTTTTTGGAAAGTATGGTAAAATG
BEK90  GTTACCGTCTTTTTGGAAAGTATGGTAAAATG
Cons   GTTACCGTCTTTTTGGAAAGTATGGTAAAATG

           +1
WT     CAATACATACTCGTTCGTTTCAAGAATCATTT
BEK90  CAATACATACTCGTTCGTTTCAAGAATCATTT
Cons   CAATACATACTCGTTCGTTTCAAGAATCATTT

WT     TCTTTTGCTCTGCCGGTAAAAGAAAAAGTGAG
BEK90  TCTTTTGCTCTGCCGGTAAAAGAAAAAGTGAG
Cons   TCTTTTGCTCTGCCGGTAAAAGAAAAAGTGAG

                  SD          CDS
WT     ACAAGATTAAGGGGTGAAAGAATG
BEK90  ACAAGATTAAGGGGTGAAAGAATG
Cons   ACAAGATTAAGGGGTGAAAGAATG
```

```
                Protein alignment

                 10            20            30
WT     MFKFNEEKGQLKCSFCGKTQDQVRKLVAGPG
BEK90  MFKFNEEKGQLKCSFCGIRRKEEGNNKWLK-
Cons   MFKFNEEKGQLKCSFCG  #/#  #
                   Zn finger
                    motif A
```

**Supplementary Figure S2: BEK90 *clpX* strain contains remaining truncated ClpX**

DNA (left) and protein (right) alignment of a *B. subtilis* wild-type and the *clpX* mutant (BEK90) from Gerth *et al.* (1). Boldface type display sequence identity between the wild-type and the BEK90 strain. The -35 and -10 boxes correspond to the SigA promoter of the *clpX* gene, +1 represents the transcriptional start site, SD the Shine Dalgarno and CDS the start of the coding sequence. Within the protein alignment of the native ClpX and the residual N-terminal ClpX of BEK90, the # indicates conserved residues and / semi-conserved residues.

**Supplementary table S1.** Primers used for strain generation and NIR Northern blot probes

| Purpose | Name | Sequence (5'-3') |
|---|---|---|
| BZZ01 | DclpC_up_for | CCGGTGATATTCTCATTTTAGTCATATCGATTCATCCTCCTG |
| | DclpC_up_rev | CCGGTGATATTCTCATTTTAGTCATATCGATTCATCCTCCTG |
| | DclpC_do_for | CATCTTACTCGATGAACTCTTCTAATATAGAAGACGGAAATGAGGC |
| | DclpC_do_rev | CTATTCCCGTTGCCATACGCC |
| BZZ02 | DclpE_up_for | CTCGTAATATAAGCCGTGATC |
| | DclpE_up_rev | ATATAAGCAATCAGTCATGTCAAGTCTTATGCACAAAAATTTTTG |
| | Res_SigA_Phleo_for | TTGACATGACTGATTGCTTATATTATAATGTCAAAGTAACACGAAGGAGGAGGG |
| | | TAAGATGTTACAGTCTATCCCGGC |
| | Res_Phleo_just_rev | TTAGCTCTTGATCTGTTGGAAG |
| | DclpE_do_for | CTTCCAACAGATCAAGAGCTAACAAGCTGCTAATTTCAGTAGACC |
| | DclpE_do_rev | GAGAAGAGTAAGGATGTCGG |
| BZZ03 | DclpP_up_for | TCCATCGGAACAGGTGAAGC |
| | DclpP_up_rev | CATAATGCTCCTCCTTCACC |
| | Res_ Spec_for | GGTGAAGGAGGAGCATTATGAACACGTACGAGCAGATC |
| | Res_Spec_rev | GTCAATCAGGCCGTATTCAAGTTACAACTTCTTTAAGCGGTTGTTC |
| | DclpP_do_for | CTTGAATACGGCCTGATTGAC |
| | DclpP_do_rev | CTGTTGCAGGAGAATGATCC |
| BZZ04 | DclpX_up_for | CATCCCTGGCTTCGAAGATC |
| | DclpX_up_rev | TTTGTTCATTCTTTCACCCCTTAATCTTGTCTCACTTTTTCTTTTACCGG |
| | Res_clpX_Ery_for | GGGGTGAAAGAATGAACAAAAATATAAAATATTCTCAAAAC |
| | Res_clpX_Ery_rev | TTATTTCCTCCCGTTAAATAATAG |
| | DclpX_do_for | CTATTATTTAACGGGAGGAAATAAAGATAAGCACAAACCTCCTGAG |
| | DclpX_do_rev | CAACCGTTTTCAGCTCGTCC |
| BZZ05 | DctsR_up_for | CATTCGCAGAGTGTAAAGGC |
| | DctsR_up_rev | GGTGATATTCTCATTTTAGTCATTCAACCCCCTCCTTTACTG |
| | Res_Km_just_for | ATGACTAAAATGAGAATATCACCGG |
| | Res_Km_just_rev | GAAGAGTTCATCGAGTAAGATGTAGTACTTGATTTTCTCCCAATCAGGCTTGATCC |
| | DctsR_do_for | CTTACTCGATGAACTCTTCTAAGCGGGTGAAAAGATTGATTTG |
| | DctsR_do_rev | CTTCAATCCAGTCGTCGACC |
| BCSH01 | 1_cond_ClpX_up_for | AATCGAGCCTGTAGACCGTCCTG |
| | 2_cond_ClpX_up_rev | ACGTTTGCGTGCCAATTCGTTTCTGACTTGCACATTCTATATG |
| | 3_TetR_PsigA-TRE_for | GAATTGGCACGCAAACGTAAAC |
| | 4_TetR_PsigA-TRE_rev | AAGGATCCCTATCACTGATAGGGAATCTATCTTAATTATATC |
| | 5_cond_ClpX_do_for | ATCAGTGATAGGGATCCTTTACTCGTTCGTTTCAAGAATC |
| | 6_cond_ClpX_do_rev | CTCCTGAGTGTTACCACTCAGGAG |
| *clpE* probe | clpE_Nor_for | GCATCTCTTGTTGCAAACAC |
| | clpE_T7_rev | GAAATTAATACGACTCACTATAGGGAGAATGATAAAGTGACACATGCTTTGATTG |

**Supplementary table S2.** Parameter for mass spectrometric analysis using an Orbitrap Exploris™ 480 mass spectrometer (Thermo Fisher Scientific) coupled to an UltiMate™ 3000 RSLC nano system (Thermo Fisher Scientific)

| | |
|---|---|
| **Reversed phase liquid chromatography (RPLC)** | |
| *Instrument* | Ultimate 3000 RSLC (Thermo Fisher Scientific) |
| *Trap column* | 75 µm inner diameter, packed with 3 µm C18 particles (Acclaim PepMap100, Thermo Scientific) |
| *Analytical column* | Accucore 150-C18, (Thermo Fisher Scientific)<br>25 cm x 75 µm, 2,6 µm C18 particles, 150 Å pore size |
| *Buffer system* | binary buffer system consisting of 0.1% acetic acid in HPLC-grade water (buffer A) and 100% ACN in 0.1% acetic acid (buffer B) |
| *Flow rate* | 300 nl/min |
| *Gradient* | linear gradient of buffer B from 2% up to 25% |
| *Gradient duration* | 60 min |
| *Column oven temperature* | 40°C |
| **mass spectrometry** | |
| *instrument* | Orbitrap Exploris™ 480 mass spectrometer (Thermo Fisher Scientific) |
| *Electrospray* | Nanospray Flex Ion Source |
| *Operation mode* | data-independent |
| ***Full MS*** | |
| *MS scan resolution* | 120000 |
| *Norm. AGC target (%)* | 300 |
| *maximum ion injection time for the MS scan* | 60 ms |
| *Scan range* | 350 to 1200 m/z |
| *RF Lens* | 50 % |
| *Spectra data type* | profile |
| ***dd-MS2*** | |
| *Precursor mass range* | 350 to 1200 m/z |
| *Resolution* | 30,000 |
| *Norm. MS/MS AGC target (%)* | 3000 |
| *Maximum ion injection time mode* | auto |

| | |
|---|---|
| *Spectra data type* | profile |
| *Microscans* | 1 |
| *Isolation window* | 66 windows, 13 m/z, 2 m/z overlap |
| *Define first mass* | 200 |
| *Dissociation mode* | higher energy collisional dissociation (HCD) |
| *HCD normalized collision energy* | 30% |
| *Samples excluded after measurement based on principle component analysis* | clpC $t_0$ BR3, clpP $t_0$ BR1, clpP $t_{10}$ BR1, clpP $t_{30}$ BR1, clpP $t_0$ BR2, clpP $t_{10}$ BR2, clpP $t_{30}$ BR2 |

**Supplementary table S3.** *Parameter for mass spectrometric analysis using Bruker TIMS TOF HT (Bruker Daltonics GmbH) coupled to an Evosep One System (Evosep Biosystems Aps)*

| Nano RPLC | |
|---|---|
| *Column system* | Evosep One (Evosep Biosystems Aps)<br>Evotip Pure, sample loaded by following guide given by Evosep<br>Evosep Performance column EV-1137, 15cm x 150um, 1.5um C18 column heated to 40degC<br>30SPD LC method by Evosep |
| *Buffer system* | binary buffer system consisting of 0.1% formic acid in HPLC-grade water (buffer A) and 100% ACN in 0.1% formic acid (buffer B) |
| **mass spectrometry** | |
| *Electrospray* | Captive spray 2 Emitter 10 um part no, 1811112, Spraying Voltage: 1700v |
| *Operation mode* | data-independent |
| *MS and MS/MS scan range* | 100-1700m/z positive polarity data acquired with dia-PASEF mode activated in 400-1001 m/z, 30 m/z windows and 1.17 sec cycle time |
| *mobility range* | 0.65-1.45 V-s/cm2 |
| *ramp and accumulation time* | 100ms |
| *collision energy ramping* | for mobility range 0.6-1.60 V-s/cm2 was 20-59eV |
| *Samples excluded after measurement based on principle component analysis* | wildtype $t_0$ BR1, wild-type $t_{30}$ BR3, *clpX* $t_{10}$ BR3 and *con clpX* ctrl BR3 |
